# Longitudinal prevalence and co-carriage of pathogens associated with nursing home acquired pneumonia in three long-term care facilities

**DOI:** 10.1101/2024.12.19.629505

**Authors:** Ryann N. Whealy, Alexander Roberts, Tara N. Furstenau, Skylar Timm, Sara Maltinsky, Sydney Wells, Kylie Drake, Kayla Ramirez, Candice Bolduc, Ann Ross, Talima Pearson, Viacheslav Y. Fofanov

**Author notes:** <u>Corresponding authors:</u> Viacheslav Y. Fofanov, 1295 Knoles Dr, Flagstaff, AZ 86011, USA, Talima Pearson, 1395 Knoles Dr, Flagstaff, AZ 86011, USA.

## Abstract

Nursing home acquired pneumonia (NHAP), and its subset – aspiration-associated pneumonia, is a leading cause of morbidity and mortality among residents in long-term care facilities (LTCFs). Understanding colonization dynamics of respiratory pathogens in LTCF residents is essential for effective infection control. This study examines the longitudinal trends in prevalence, persistence, bacterial load, and co-colonization patterns of five respiratory pathogens in three LTCFs in Phoenix, Arizona. Anterior nares and oral swabs were collected every other week and tested using qPCR for *Haemophilus influenzae*, *Pseudomonas aeruginosa*, *Streptococcus pneumoniae*, *Staphylococcus aureus*, and *Chlamydia pneumoniae*. Weekly average positivity rates were 17.75% for *H. influenzae* (0% – 39.39%), 9.95% for *P. aeruginosa* (0% – 37.74%), 31.89% for *S. pneumoniae* (1.79% – 41.67%), and for 28.00% for *S. aureus* (0% – 55.36%). *C. pneumoniae* was not detected. *H. influenzae* and *S. pneumoniae* predominantly colonized the oral cavity, while *P. aeruginosa* and *S. aureus* predominantly colonized the nasal cavity. *S. pneumoniae* and *S. aureus* colonizations were significantly more persistent than *H. influenzae* and *P. aeruginosa*, with persistence correlating with significantly higher bacterial loads. Co-colonization did occur in ∼20% of positive samples, but appeared to be due to random chance. This study reveals distinct colonization patterns among respiratory pathogens in LTCF residents, highlighting differences in site-specific prevalence, persistence, and bacterial load. These findings underscore the importance of longitudinal monitoring to inform targeted infection control strategies in LTCFs.

## Introduction

Nursing home acquired pneumonia (NHAP) is a leading cause of morbidity and mortality in long term care facilities (LTCFs), accounting for 40% of hospitalizations and up to 33% of deaths among residents [1]. It is estimated that 365 out of every 1000 LTCF residents are diagnosed with pneumonia annually [2], a rate 6-11 times higher than the incidence of pneumonia in elderly people living in the community [3,4]. Further, the medical care and treatment required for NHAP contribute to significant healthcare costs due to the potential for prolonged hospital stays, need for intensive care, and treatment-resistant etiologies. While pneumonia can often be treated by LTCF staff at an average cost of approximately $3,700 [5], up to 30% of patients require hospitalization which increases average treatment cost to $10,400 [6,7]. For patients who require hospitalization, mortality rates range from 13-45% [8,9]. These high costs and poor outcomes underscore the need for targeted, evidence-based infection prevention and stewardship strategies in nursing homes [10].

Micro-aspiration is the primary pathogenic mechanism of pneumonia in otherwise healthy individuals, while macro-aspiration (the cause of aspiration pneumonia) typically occurs only in those experiencing difficulty swallowing or changes in consciousness [11]. Older individuals are more likely to sustain an infection induced through aspirating large amounts of opportunistic pathogens due to decreased humoral and cell-mediated immunity, increased likelihood of underlying airway disorders, and altered levels of consciousness. Many bacterial etiological agents of pneumonia asymptomatically colonize the oropharyngeal mucosa or other areas of the body. As such, sources of many pneumonia infections are endogenous and due to autoinfection [12]. Controlling pneumonia from endogenous sources requires management and control of these pathogens at individual and institutional levels to reduce the likelihood of acquisition, carriage, and infection.

Many pneumonia-causing bacteria colonize dental plaque biofilms [13–16], providing a potential source for micro– and macro-aspiration pneumonia [14,16]. Up to 80% correlation has been found between aerobic dental plaque colonization and the causative agent of aspiration associated-pneumonia [16]. Unsurprisingly, poor oral hygiene increases the risk of pneumonia [14,17–19], highlighting its potential as an intervention target. Further, elderly LTCF residents tend to have particularly poor oral health with high plaque scores [20], indicating that improving oral health among this population may be especially impactful. While promising, oral health interventions have shown mixed results in lowering NHAP incidence. In some studies, improved oral hygiene has been shown to decrease the incidence rate of pneumonia, potentially preventing as many as 10% of NHAP deaths [16,21–24]. Other studies have found no advantage in toothbrushing and chlorhexidine rinses over standard care in preventing NHAP [25]. In short, while the relationship between colonization and infection is well established, much research still needs to be done to assess the potential for oral hygiene interventions in reducing NHAP.

Due to the varied etiological agents involved in NHAP, characterization of longitudinal oral colonization and associated transmission dynamics are critical for guiding future intervention studies. Furthering the understanding of the transmission dynamics of the most common etiological agents responsible for NHAP (and, simultaneously, developing better assays to detect them) will enable future intervention studies aimed at equipping LTCF clinicians with strategies for mitigation and prevention. Towards this end, this study characterizes the colonization dynamics of common etiological agents of NHAP in the oral and nasal cavity in LTCF residents.

## Methods

### Sample and data collection

Across three LTCFs in the Phoenix metropolitan area, 121 residents were enrolled to participate in this study after providing written consent (IRB: 1766728). Participants provided their pneumonia vaccination status, as well as demographic information (age, sex, and ethnicity). Swabs of the anterior nares and oral mucosa were collected every other week from November 2021 to November 2023 (49 events total). Oral hygiene professionals supervised the self-sampling, which has proven to be an effective and sensitive method of detecting pathogens at these body sites [26]. Collected swabs were stored in 1mL of Liquid Amies at –20°C until processing.

### Sample processing, testing, and sequencing

#### Multi-pathogen qPCR panel

Total DNA was extracted from nasal and oral swab samples using the Applied Biosystems MagMAX Viral and Pathogen Nucleic Acid Isolation Kit with the KingFisher Flex system, according to the manufacturer’s instructions. qPCR was performed using a multi-pathogen panel (adapted from published assays) [27–31] designed to detect and quantify *Haemophilus influenzae*, *Chlamydia pneumoniae*, *Pseudomonas aeruginosa*, *Streptococcus pneumoniae*, and *Staphylococcus aureus* (Supplemental Table 1). We performed an in-silico PCR to evaluate the specificity of the primers and probes across a comprehensive set of reference genomes for each pathogen. Some oligonucleotide sequences were adjusted to maximize complementarity with a wider range of genomes. The assays were validated independently using positive controls, and optimized to be run in duplex. There were interactions between oligos for *H. influenzae* and *S.* pneumoniae as well as *S. aureus* and *P. aeruginosa*, but no other combinations produced cross-primer dimerization based on in-silico PCR. Serial dilutions of quantitative genomic DNA were included in all runs to allow for relative DNA quantification using standard curve analysis.

Each qPCR reaction was carried out in a 10 μL volume containing 5 μL of TaqMan Universal Master Mix, 2 μL of template DNA, final concentrations of 0.6 μM for primers and 0.3 μM for probes, and nuclease-free water. Cycling conditions were as follows: 50°C for 2 minutes, 95°C for 15 minutes, then 45 cycles of 95°C for 15 seconds and 57°C for 1 minute. Amplifications were completed using an Applied Biosystems QuantStudio™ 7 Pro System, and results were analyzed using the QuantStudio™ Design & Analysis Software (version 2.6.0).

#### Secondary validation using multi-pathogen amplicon sequencing

To confirm that the multiplex qPCR assays accurately identified the target species, we used a species-specific targeted amplicon sequencing approach to detect additional genomic regions unique to each pathogen in a subset of qPCR positive samples (Supplemental Table 2). The multiplex PCR was performed using the KAPA 2G Fast Multiplex PCR Master Mix in 25µL reaction volumes, with 5ng of template DNA, 0.2 µM concentration of each primer, and water to reach the final volume. PCR thermocycling conditions consisted of an initial denaturation at 95°C for 3 minutes, followed by 35 cycles of denaturation at 95°C for 15 seconds, annealing at 60°C for 30 seconds, and extension at 72°C for 1 minute and 30 seconds and a final extension at 72°C for 1 minute. PCR products were prepared for sequencing using our standard amplicon sequencing protocol [32] and were sequenced on the Illumina MiniSeq platform. Reads were aligned to reference genomes (Accessions in Supplemental Table 2) using BWA-MEM v0.7.17 [33].

### Data analysis

All data cleaning and statistical analyses were conducted using R version 4.4.1. Statistical significance was set at a p-value of less than 0.05 for all tests. To minimize bias from sampling inconsistency, results are only included from participants who attended at least ⅓ of sampling events.

McNemar’s test [34] was used to compare colonization rates between nasal and oral samples for each pathogen, accounting for the paired nature of the samples collected from the same participants. Matched odds ratios with 95% confidence intervals were calculated to quantify the magnitude of differences in colonization rates. To estimate the probabilities of transitioning between colonization states over time, we employed a Markov chain analysis with bootstrapping (1000 for each pathogen). The resulting transition matrices generated from bootstrapping were averaged to produce a final transition matrix representing the mean transition probabilities. Linear mixed models were fitted using the lme4 [35] and lmerTest [36] packages to quantify predictors of average relative DNA quantities. Full models included transient vs. persistent colonization, age, sex, pneumococcal vaccination, and recent estimated exposure as fixed effects as well as random intercepts for each participant to account for repeated measures. The results are reported in relative units (RQ), indicating fold changes in bacterial abundance relative to a standardized control. Co-colonization patterns were visualized using an upset plot created with the UpSetR [37] package, highlighting the intersections between different pathogens. To investigate potential temporal associations between colonization with different pathogens, we performed a correlation analysis incorporating current and lagged colonization statuses. However, no significant correlations were found, so this analysis was exploratory in nature.

## Results

### Participant description

The 121 participants ranged in age from 60 to 97 (median = 83, mean = 82.97) and were a majority female (76.03%). Race/ethnicity was not included in this analysis due to a significantly imbalanced distribution within the sample; approximately 96% of participants were non-Hispanic whites. The subset of participants who attended at least one-third of sampling events (n=85) closely matched the demographic distribution of the overall study population, with a mean age of 83 and approximately ¾ being female. These individuals attended an average of 34 out of 49 sampling events, approximately 70% were vaccinated against streptococcal pneumonia, and were evenly distributed between the three LTCFs.

### Colonization rates at nasal and oral sites

We observed wide ranges of colonization rates for the pathogens of interest and significant site-specificity towards the nasal or oral cavity. Of the approximately 6,000 samples collected from the 85 participants in LTCFs, 9.99% of samples were positive for *Haemophilus influenzae*, 5.78% were positive for *Pseudomonas aeruginosa*, 22.07% were positive for *Streptococcus pneumoniae*, and 18.01% were positive for *Staphylococcus aureus*. Notably, *Chlamydia pneumoniae* was not detected in any samples. Table 1 provides an overview of the colonization metrics for each pathogen. Across time points, notable variation was observed in both sample-level and individual-level positivity rates. The cumulative incidence values illustrate that a majority of participants were colonized by at least one of these pathogens during the study period.

**Table 1:** Colonization Metrics by Pathogen.

| Pathogen | Average weekly sample positivity (range) | Average weekly individual positivity (range) | Cumulative incidence |
| --- | --- | --- | --- |
| *H. influenzae* | 9.81% (0–23.48) | 17.75% (0–39.39) | 81.18% |
| *P. aeruginosa* | 5.56% (0–20) | 9.95% (0–37.74) | 83.53% |
| *S. pneumoniae* | 21.70% (0.89–31.62) | 31.89% (1.79–41.67) | 78.82% |
| *S. aureus* | 17.80% (0–36.13) | 28.00% (0–55.36) | 91.76% |

Our findings indicate that these pathogens differentially colonize the oral and nasal cavities, with *H. influenzae* and *S. pneumoniae* predominating in the oral cavity, and *P. aeruginosa* and *S. aureus* more frequently detected in the nasal cavity (Table 2). These differences were statistically significant (McNemar’s test, p < 1×10^-6), suggesting that the observed distribution patterns are not due to random variation but reflect true biological differences in site-specific colonization. Matched odds ratios further quantify these disparities, demonstrating substantially reduced odds of nasal colonization for *H. influenzae* and *S. pneumoniae* relative to oral colonization, but notably increased odds of nasal colonization for *P. aeruginosa* and *S. aureus*.

**Table 2:**
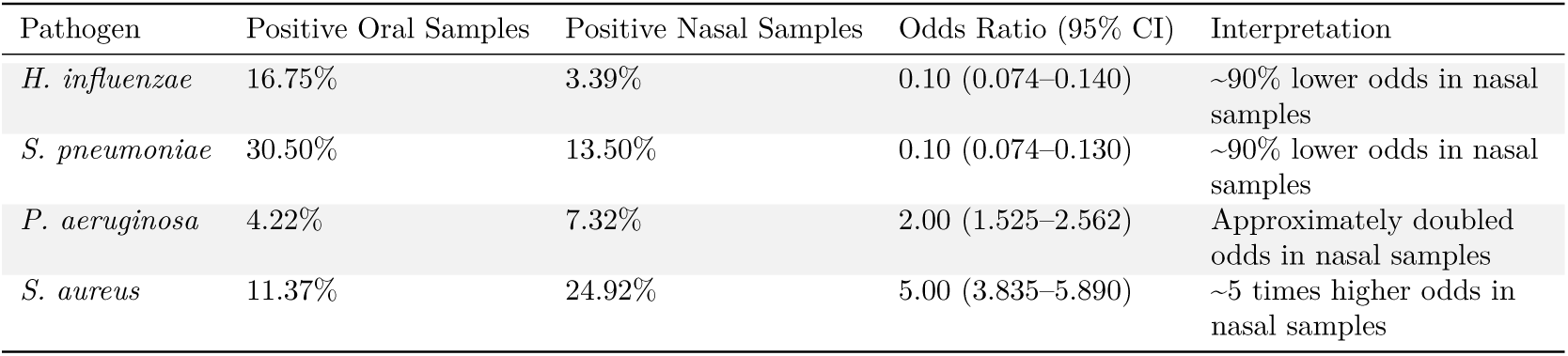
Site-Specific Colonization.

### Longitudinal colonization rates

There were statistically significant changes in participant positivity over time for some of the pathogens (Figure 1). *H. influenzae* colonization decreased over time in both sample types, while *P. aeruginosa, S. aureus,* and *S. pneumoniae* colonization decreased in nasal samples over time (p < 0.01). Oral colonization with *P. aeruginosa, S. pneumoniae,* and *S. aureus* did not significantly change over time. Colonization at the individual level changed significantly over time for *H. influenzae, P. aeruginosa,* and *S. aureus* (p < 0.01). These changes were not correlated with any measured variables and there were no changes in protocol during the study.

**Figure 1.**
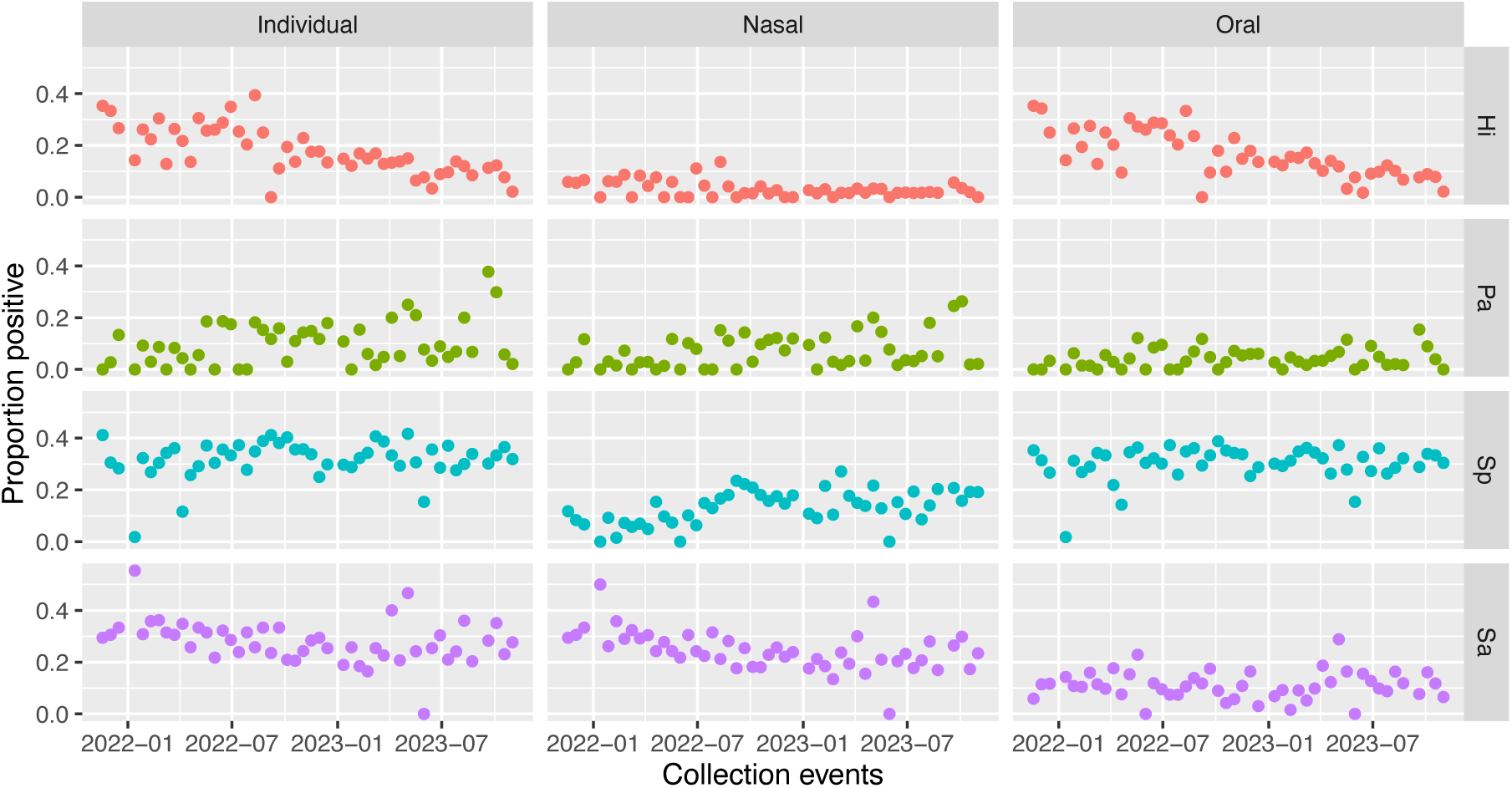
The proportion of individuals (from oral, nasal, or both sample types), nasal samples, and oral samples testing positive for each pathogen over the course of the study. Pathogens include *Haemophilus influenzae* (Hi, red), *Pseudomonas aeruginosa* (Pa, green), *Streptococcus pneumoniae* (Sp, blue), and *Staphylococcus aureus* (Sa, purple). Each dot represents the positivity rate for a collection event, with trends highlighting temporal dynamics in colonization prevalence across anatomical sites and pathogens.

### Duration of colonization

The distribution of time participants spent colonized shows notable variability among pathogens (Figure 2). The proportion of time that participants spent colonized by *S. pneumoniae* and *S. aureus* exhibited a pronounced bimodal pattern with a peak near zero, indicating never or highly transient colonization, and another near one, reflecting persistent colonization. Interestingly, colonization by *P. aeruginosa* appears to be predominantly short-lived, with only rare cases of persistent colonization or carriage. Colonization with *H. influenzae* was similarly skewed, though less drastically. The observation that individuals tend to be colonized by *S. aureus* either a majority of the time or only transiently has previously been found [38,39]. Less is known about persistence of colonization with the other pathogens of interest in healthy individuals. To quantify this variability, participants were categorized as “never” (no colonization over the study), “transient” (colonization in <50% of sampling events), or “persistent” (colonization in ≥50% of sampling events). For *H. influenzae*, 67.1% of participants experienced transient colonization, 18.8% were never colonized, and 14.1% were persistently colonized, indicating that *H. influenzae* colonization is mostly short-lived for otherwise healthy individuals.

**Figure 2.**
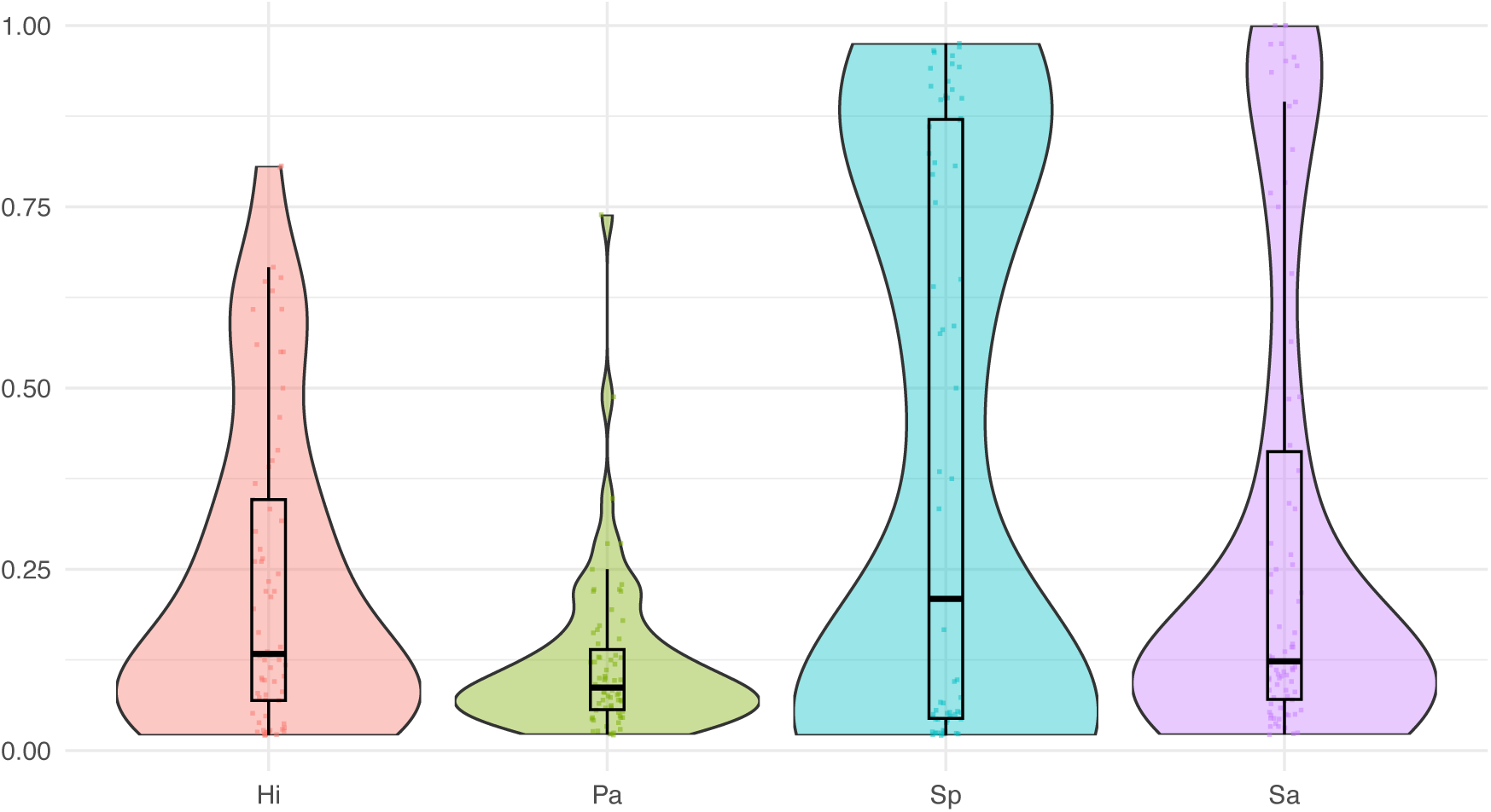
The proportion of events participants spent colonized by each pathogen (if colonized at least once); *Haemophilus influenzae* (Hi, red), *Pseudomonas aeruginosa* (Pa, green), *Streptococcus pneumoniae* (Sp, blue), and *Staphylococcus aureus* (Sa, purple). The distribution indicates variability in colonization duration across pathogens, with box plots embedded to show median and quartile ranges.

For *P. aeruginosa*, 82.4% were transient, 16.5% were never colonized, and 1.18% were persistently colonized, indicating that *P. aeruginosa* is not typically a long-term colonizer. In contrast, *S. pneumoniae* exhibited a more balanced pattern: 43.5% were transiently colonized, 21.2% were never colonized, and 35.3% were persistently colonized. For *S. aureus*, only 8.24% were never colonized, with 71.8% transiently colonized and 20% persistent colonization.

The average duration of continuous colonization at any of the two body sites was approximately 4 weeks for *H. influenzae*, 2.4 weeks for *P. aeruginosa*, 9.6 weeks for *S. pneumoniae*, and 8.8 weeks for *S. aureus* (of those that were colonized at least once during the study). Recurrent colonization (recolonization after pathogen was not detected and thus assumed to be cleared from an individual) was observed in 58.8% of participants for *H. influenzae*, 61.2% for *P. aeruginosa*, 57.6% for *S. pneumoniae*, and 72.9% for *S. aureus*. These colonization patterns are illustrated below (Figure 3). *S. pneumoniae* and *S. aureus* are often persistent colonizers, while *H. influenzae* and *P. aeruginosa* typically appear transiently. Interestingly, persistent colonization with *S. aureus* tends to be in the nasal cavity while persistent colonization with *S. pneumoniae* tends to occur in the oral cavity (Figure 3).

**Figure 3.**
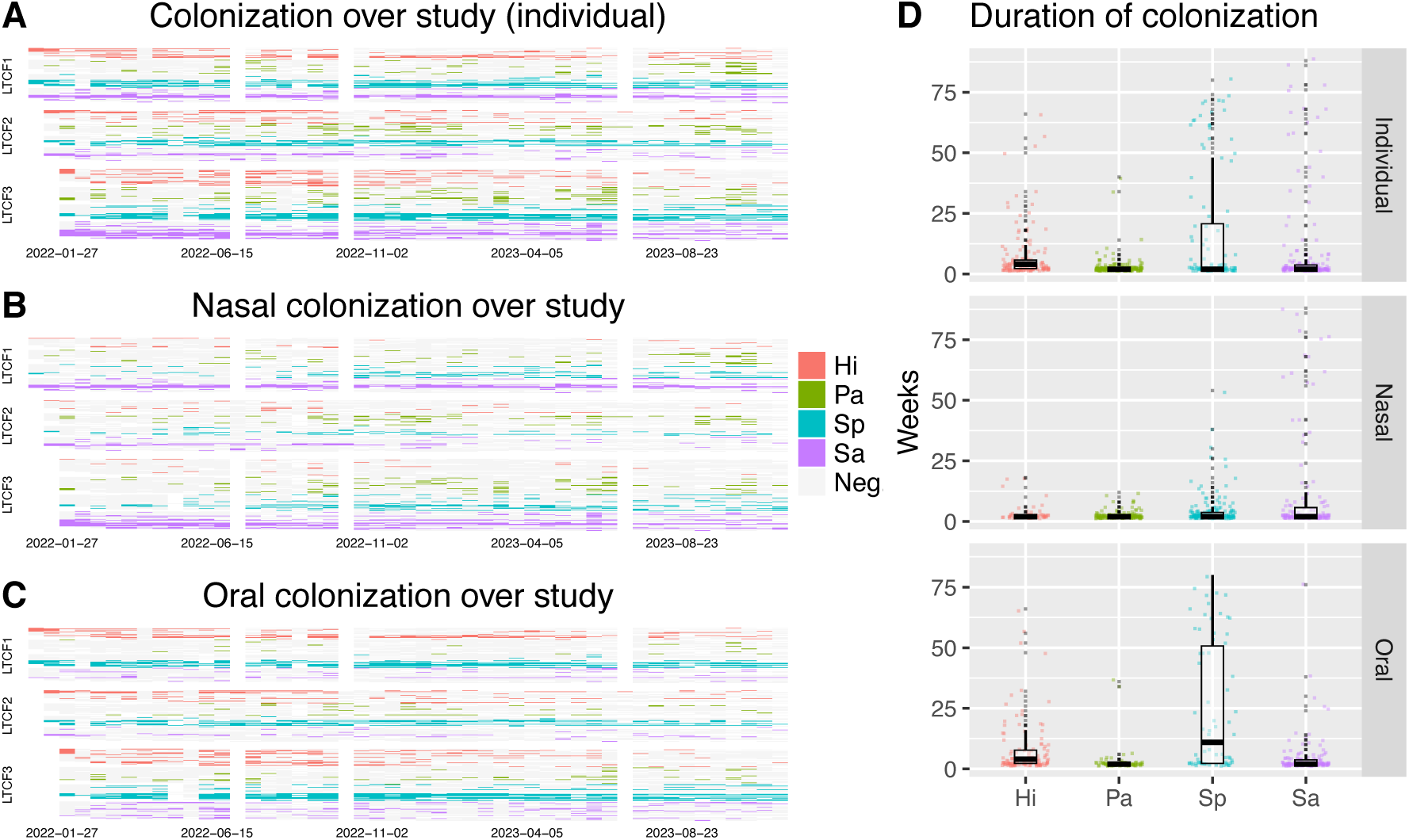
Colonization patterns of *Haemophilus influenzae* (Hi, red), *Pseudomonas aeruginosa* (Pa, green), *Streptococcus pneumoniae* (Sp, blue), and *Staphylococcus aureus*. (Sa, purple) shown by A) Colonization at the individual level (from oral, nasal, or both sample types) across the study period where each row represents an individual participant grouped by their LTCF; B) Nasal colonization across the study period, showing varying persistence of pathogens over time in each sampled LTCF; C) Oral colonization across the study period, showing varying persistence of pathogens over time in each sampled LTCF; and D) Duration of continuous colonization for each pathogen at the individual level and at nasal and oral sites, highlighting variability.

Markov chains were used to provide a more nuanced understanding of colonization likelihood and stability over time. For *H. influenzae*, at any given time point, the probability of becoming colonized was 9.05%, with a 55.92% chance of maintaining colonization once established, indicating moderate persistence. *P. aeruginosa* showed a similar colonization probability of 9.05%, but only a 21.67% probability of remaining colonized, reflecting its low likelihood of sustained colonization. In contrast, *S. pneumoniae* had a 7.56% probability of colonizing but an 83.64% chance of remaining colonized, suggesting that *S. pneumoniae* is more persistent once established. For *S. aureus*, the colonization probability was 10.74%, with a 72.98% likelihood of continuing colonization once established.

To identify which factors most influenced bacterial load (and thus the amount of bacterial material available for macro-aspiration), relative DNA quantities during positive weeks were analyzed using linear mixed models for each pathogen. Participants who were never colonized by the pathogen of interest were excluded from this analysis to focus on the comparison between transient and persistent colonization. For all pathogens, a subjects’ categorization of persistent or transient colonization was the most significant predictor of their bacterial load (except for *P. aeruginosa* which did not have a pronounced persistent category). For *H. influenzae*, *S. pneumoniae* and *S. aureus*, persistent colonization was associated with significantly higher bacterial DNA levels compared to transient colonization (p < 0.001).

### Co-colonization

While colonization by a single pathogen was most common, co-positivity was observed nearly ⅓ of the time that an individual was colonized (Figure 4). However, the co-colonization rates did not appear to be more frequent than what is expected by chance (p > 0.4, under assumption of independence of colonization events). Additionally, lagged colonization status analysis suggested that there is no evidence that colonization by one pathogen predicts colonization by another within the next month (r < 0.2).

**Figure 4.**
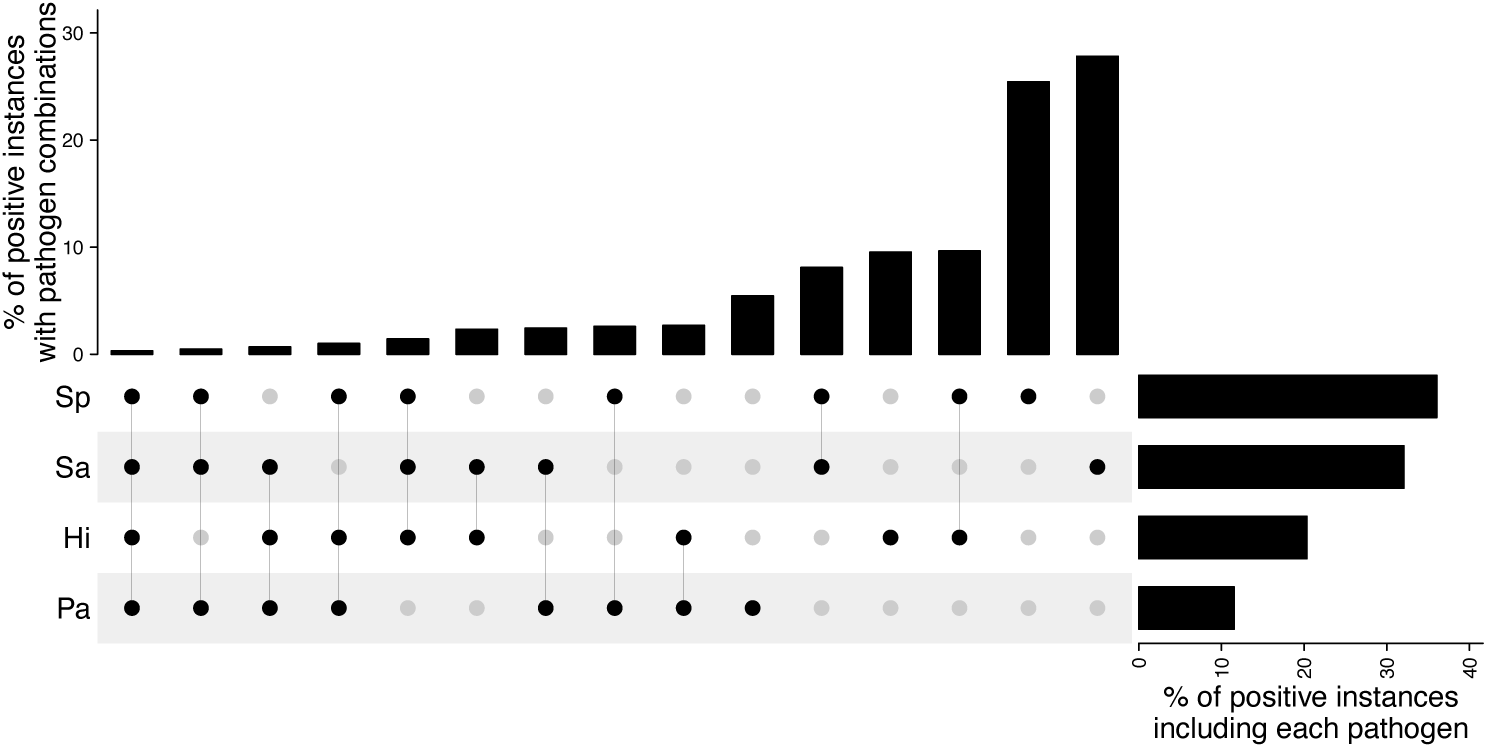
Co-colonization observed throughout the study within individuals (i.e. considering both oral and nasal results). The bar chart at top shows the percent of instances where individuals who were positive for at least one pathogen were positive for a given combination of pathogens (filled black dots). Each row represents a pathogen, and the bar chart to the right shows the relative prevalence for each pathogen among individuals that tested positive for at least one pathogen.

## Discussion

Findings from this study highlight several important colonization patterns among LTCF residents, all of which underscore the value of longitudinal data in revealing temporal variability and individual-level dynamics that would be impossible to capture with cross-sectional approaches. This study is strengthened by sample collection from multiple body sites, investigation of multiple pathogens, and the utilization of qPCR for pathogen detection, which has been shown to be an especially reliable method of detection in older populations [40]. Longitudinal data collection provides valuable information on how prevalence, persistence, bacterial load, and co-colonization patterns are affected temporally and on an individual and institutional basis. Additionally, this study provides valuable information about asymptomatic carriage in otherwise healthy individuals; the very few existing longitudinal NHAP research studies have been limited to hospitalized patients. These insights can guide the design of effective mitigation strategies for NHAP and associated etiological agents.

While average weekly prevalence rates of *H. influenzae* (average 17.75%), *P. aeruginosa* (9.95%), *S. pneumoniae* (31.89%), and *S. aureus* (28.00%) offer valuable cross-sectional snapshots, the high cumulative incidences over the study period (81.18%, 83.53%, 78.82%, and 91.76%, respectively), and the duration of individual carriage, reveal a much broader picture of participant exposure and colonization dynamics. Without tracking these pathogens over time, the episodic nature of colonization might obscure the extent of overall exposure. For instance, while weekly rates fluctuate significantly (e.g., *S. pneumoniae* ranging from 0.89% to 31.62% and *S. aureus* from 0% to 36.13%), the cumulative incidences highlight that almost all participants were colonized at least once over the course of a two-year period. These findings emphasize that point-prevalence measures alone underestimate the true burden of colonization, potentially leading to an incomplete understanding of pathogen carriage, transmission, and risks.

This study demonstrates distinct colonization patterns of these pathogens in the oral and nasal cavities. Interestingly, *Chlamydia pneumoniae* was absent from all samples, possibly due to its obligate intracellular life cycle and tendency to colonize lower respiratory tract tissue [41]. Over the two-year observation period, *H. influenzae* and *S. aureus* colonization decreased over time in their predominant niches. In contrast, nasal colonization rates of *P. aeruginosa* and *S. pneumoniae* increased as the study progressed. These temporal shifts occurred independently of participant demographics, sampling methods, or changes in study protocol, suggesting that underlying biological or ecological factors, rather than measured external variables, influenced the observed trends. While sample collection from the nasal and oral cavity allows characterization of site-specificity in these pathogens, it is likely that different strains may exhibit distinct site-specificity. Future studies may benefit from increased genotypic resolution and expanding the number of body sites sampled to further understand site-specificity and potential reservoirs for colonization and spread. Studies have shown that increasing the number of body sites sampled, increasing the number of replicates tested, and using multiple detection methods can minimize false negatives and increase prevalence estimates [26]. Consequently, it is likely that the estimates presented here (using only qPCR for detection and only one replicate from two body sites) are conservative and under-represent the true prevalence of colonization and co-colonization.

The categorization of participants into transient and persistent colonization groups provide insight into the different strategies utilized by these pathogens. Markov chain simulations confirmed the dominance of transient colonization for *P. aeruginosa,* and to a lesser degree *H. influenzae*, indicating that these pathogens frequently establish and then are cleared without long-term colonization. Analysis of bacterial loads indicated that persistent colonization is associated with a significantly higher bacterial burden for *H. influenzae*, *S. pneumoniae*, and *S. aureus*. This finding suggests that once these pathogens establish persistent colonization, they are able to maintain higher bacterial densities, which could have implications for transmission potential and clinical outcomes. This should also guide potential intervention strategies – for example, attempting to disrupt persistent *S. aureus* colonization via oral healthcare interventions (when persistent colonization appears to be in the nasal cavity) are likely to be ineffective.

Polymicrobial pneumonia represents up to 40% of cases with an identified cause [42]. The involvement of multiple pathogens can significantly affect infection dynamics, often resulting in more complex clinical outcomes [42]. In our study, co-colonization was a relatively frequent occurrence, observed approximately ⅓ of the time that an individual was colonized. However, these rates aligned with simulated expectations (p > 0.4), supporting the assumption of independence. Consistent with this, no evidence was found that colonization by one pathogen predicts subsequent colonization by another. Our findings suggest that effective preventions should focus on control of individual pathogen colonization rather than a specialized approach to multiple pathogens.

This study highlights the complex dynamics of colonization in LTCFs, including variability in site-specific colonization, the distinction between transient and persistent colonization, and the occurrence of co-colonization. Longitudinal data collection was crucial, enabling a better understanding of colonization dynamics and persistence. These insights emphasize the importance of targeted surveillance and intervention strategies that consider the temporal and specific nature of colonization, particularly for high-risk pathogens like *S. aureus* and *P. aeruginosa*.

## Author Contributions

This study was conceptualized by RNW, VYF, TP, and TNF. Data collection was carried out by CB, AR, TP, and TNF, and formal analyses were performed by RNW and TNF. Writing was completed by RNW, TNF, TP, and AR. Wet lab work and sample processing were performed by RNW, AR, ST, SM, SW, KD, and KR. All authors have reviewed the manuscript.

## Acknowledgements

This work was supported by the Centers for Disease Control and Prevention (contract 75D30121C11191 – TP) and the National Institutes of Health (NIH) National Institute on Minority Health and Health Disparities (U54MD012388 – TP) and National Institute of Allergy and Infectious Diseases (R15AI156771 – TP).

## Appendix

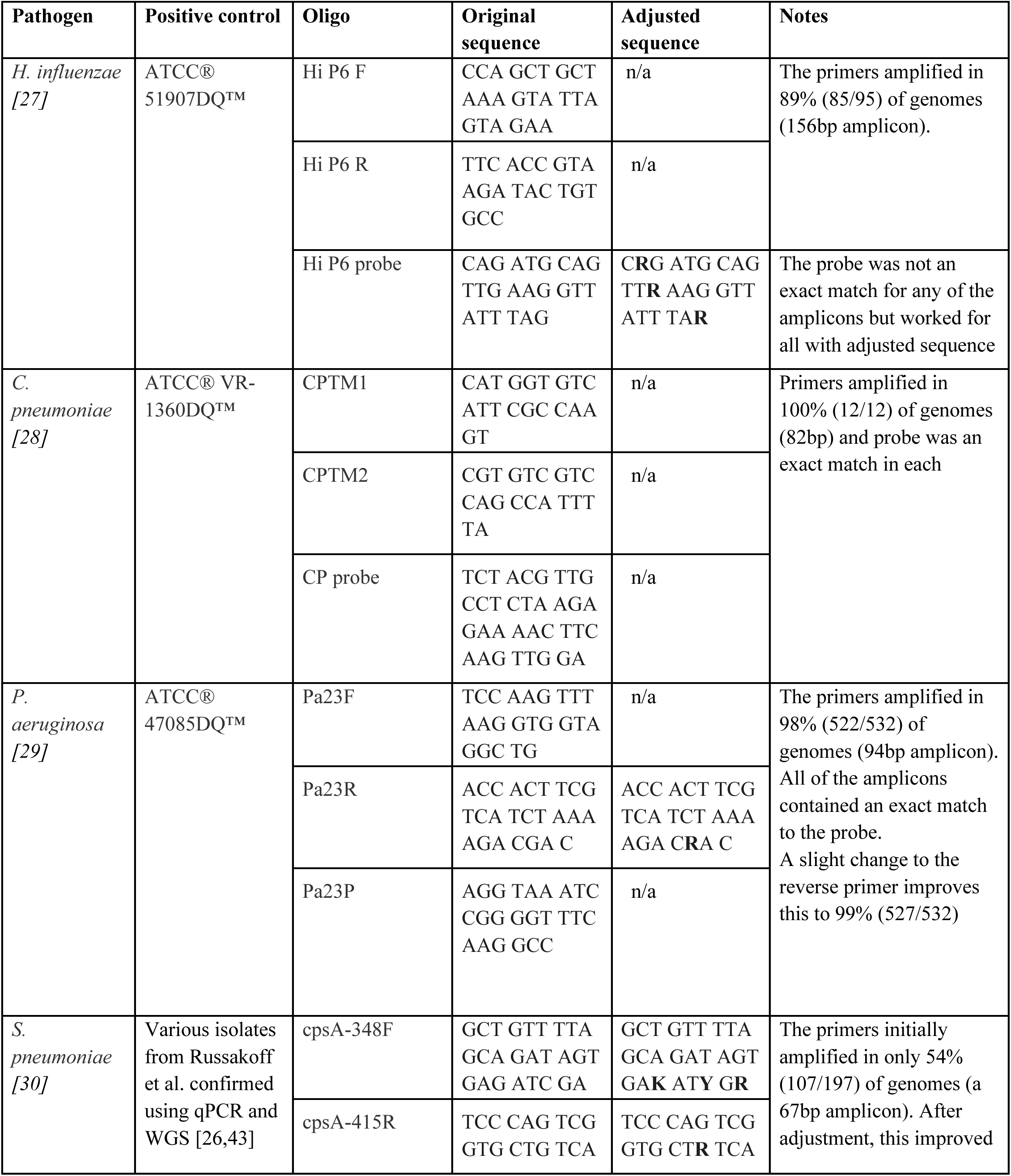
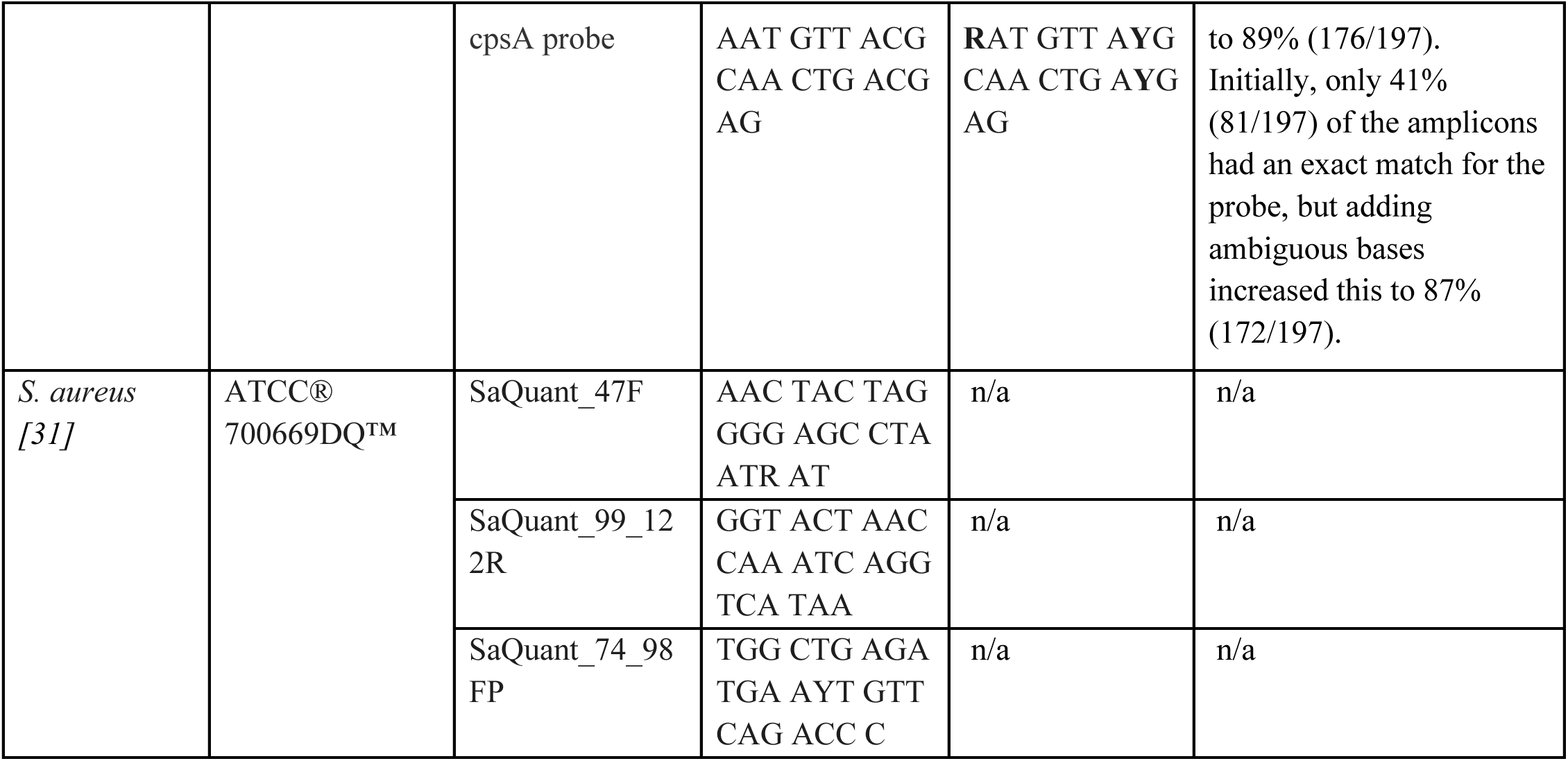
Supplemental Table 1.

**Supplemental Table 2.**
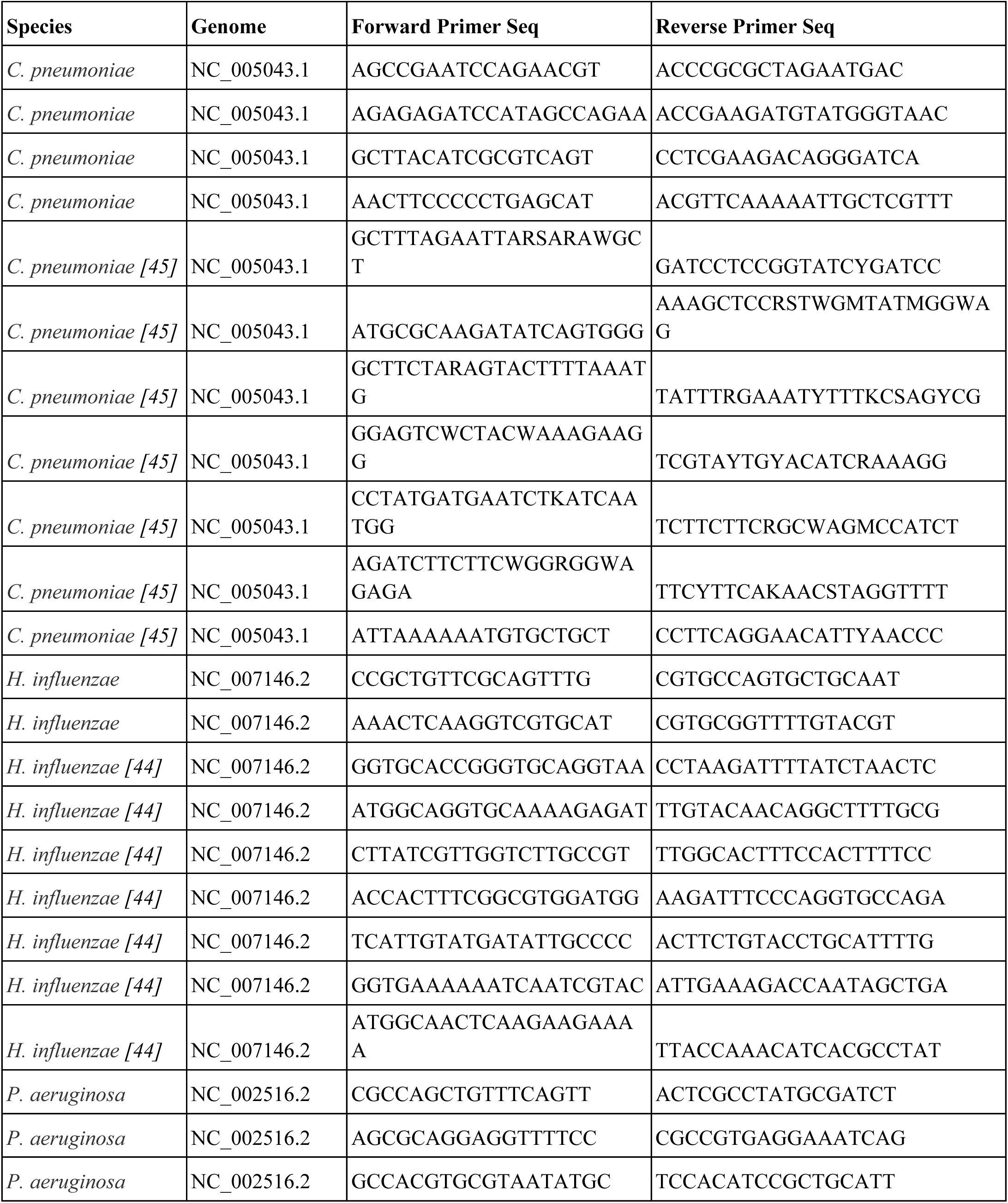
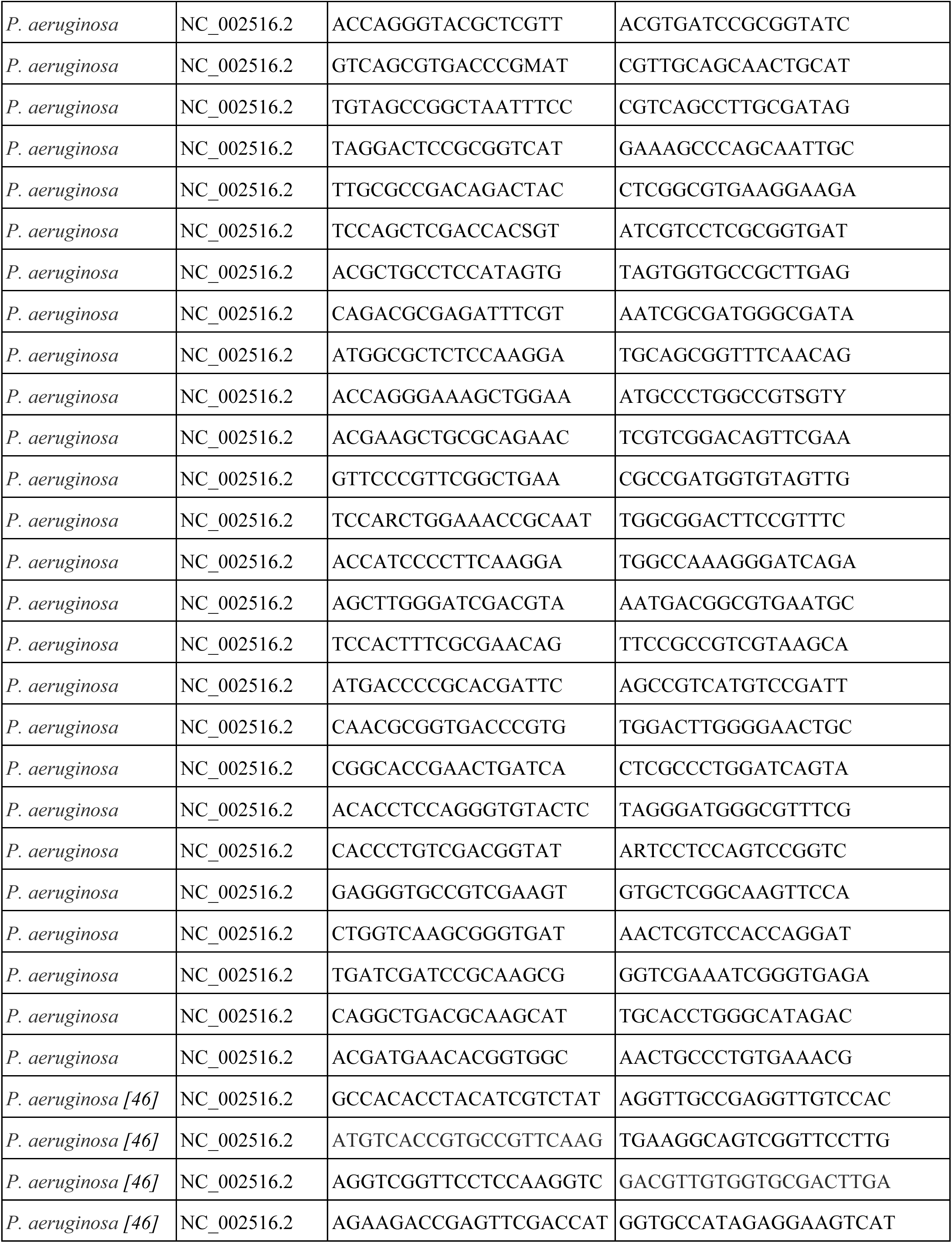
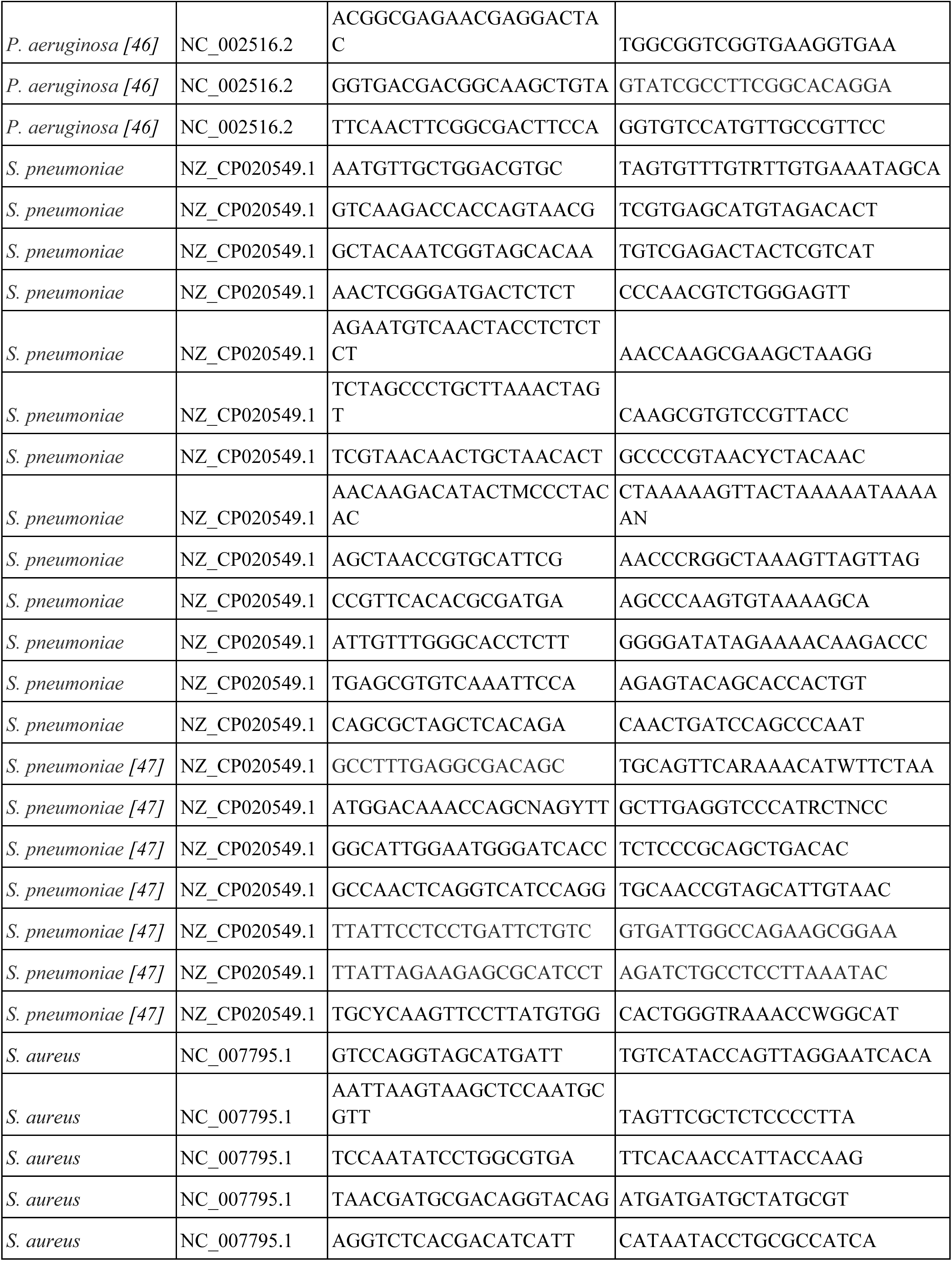
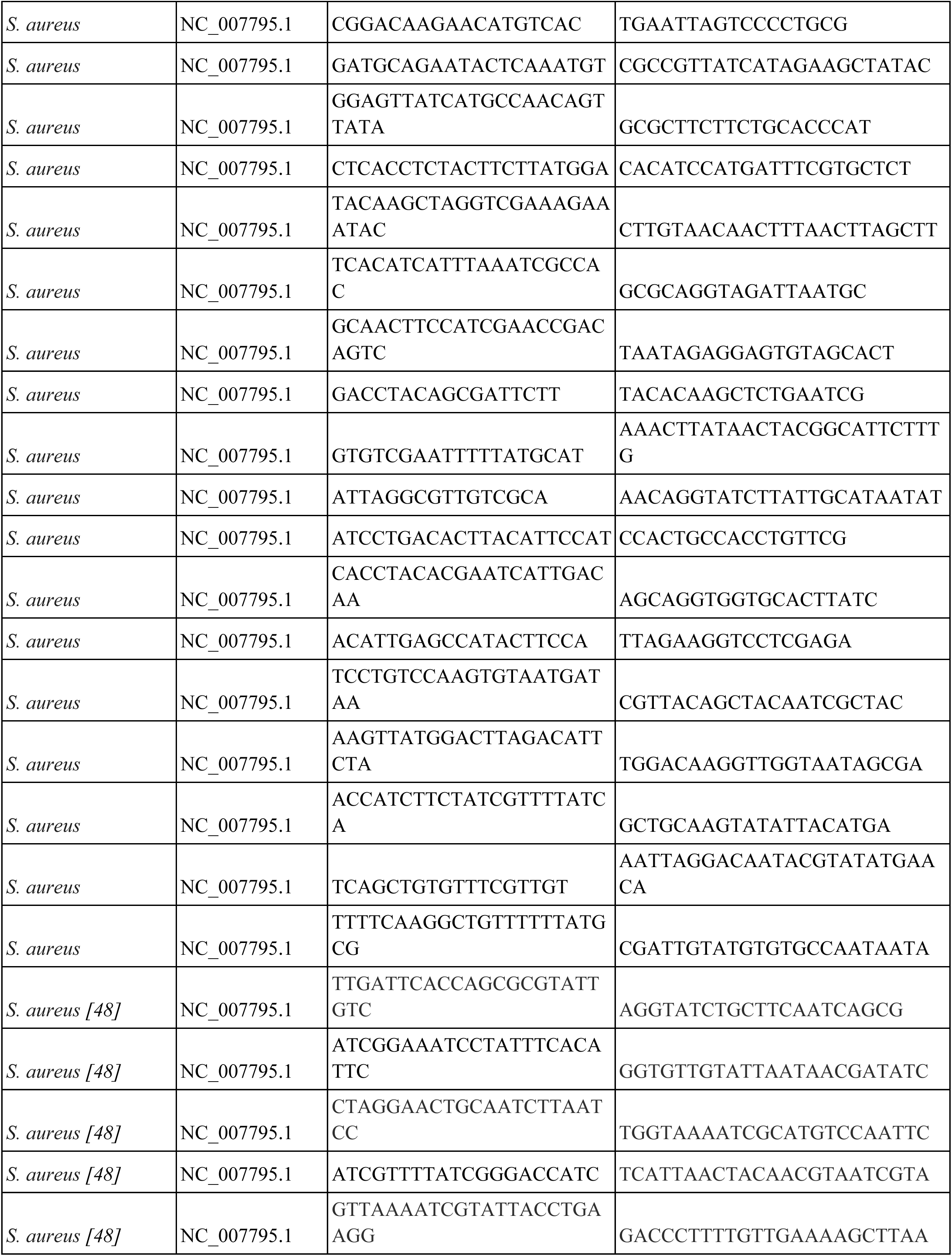
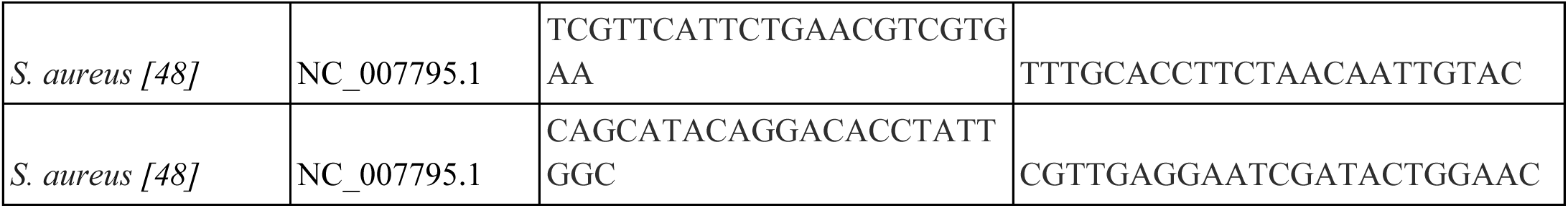
The genomic targets included established MLST targets [44–48] as well as additional pathogen-specific targets identified by selecting conserved regions from reference genome alignments within each species [49]. There were a total of 106 primer pairs: 11 for *C. pneumoniae,* 9 for *H. influenzae,* 36 for *P. aeruginosa,* 30 for *S. aureus,* and 20 for *S. pneumoniae*.

